# The Sde Phosphoribosyl-Linked Ubiquitin Transferases protect the *Legionella pneumophila* vacuole from degradation by the host

**DOI:** 10.1101/2023.03.19.533379

**Authors:** Seongok Kim, Ralph R. Isberg

## Abstract

*Legionella pneumophila* grows intracellularly within a host membrane-bound vacuole that is formed in response to a bacterial type IV secretion system (T4SS). T4SS translocated Sde proteins promote phosphoribosyl-linked ubiquitination of endoplasmic reticulum protein Rtn4, but the role played by this modification is obscure due to lack of clear growth defects of mutants. To identify the steps in vacuole biogenesis promoted by these proteins, mutations were identified that unmasked growth defects in Δ*sde* strains. Mutations in the *sdhA*, *ridL* and *legA3* genes aggravated the Δ*sde* fitness defect, resulting in disruption of the *Legionella*-containing vacuole (LCV) membrane within 2 hrs of bacterial contact with host cells. Depletion of Rab5B and sorting nexin 1 partially bypassed loss of Sde proteins, consistent with Sde blocking early endosome and retrograde trafficking, similar to roles previously demonstrated for SdhA and RidL proteins. Sde protein protection of LCV lysis was only observed shortly after infection, presumably because Sde proteins are inactivated by the metaeffector SidJ during the course of infection. Deletion of SidJ extended the time that Sde proteins could prevent vacuole disruption, indicating that Sde proteins are negatively regulated at the posttranslational level and are limited to protecting membrane integrity at the earliest stages of replication. Transcriptional analysis was consistent with this timing model for an early point of execution of Sde protein. Therefore, Sde proteins act as temporally-regulated vacuole guards during establishment of the replication niche, possibly by constructing a physical barrier that blocks access of disruptive host compartments early during biogenesis of the LCV.

**Significance statement:** Maintaining replication compartment integrity is critical for growth of intravacuolar pathogens within host cells. By identifying genetically redundant pathways, *Legionella pneumophila* Sde proteins that promote phosphoribosyl-linked ubiquitination of target eukaryotic proteins are shown to be temporally-regulated vacuole guards, preventing replication vacuole dissolution during early stages of infection. As targeting of reticulon 4 by these proteins leads to tubular endoplasmic reticulum aggregation, Sde proteins are likely to construct a barrier that blocks access of disruptive early endosomal compartments to the replication vacuole. Our study provides a new framework for how vacuole guards function to support biogenesis of the *L. pneumophila* replicative niche.

## Introduction

*Legionella pneumophila* is an intravacuolar pathogen of amoebae that can cause pneumonic disease in susceptible human hosts (1, 2). As a causative agent of Legionnaires’ disease, infection is driven by inhalation or aspiration of contaminated water, followed by bacterial growth within alveolar macrophages (3, 4). Failure to clear infection from the lungs in the immunocompromised patient results in life-threatening disease.

Successful intracellular growth in hosts depends on the establishment of the specialized *Legionella*-containing vacuolar (LCV). Upon internalization, more than 300 different effectors are translocated through the Icm/Dot type IV secretion system (T4SS) into host cells (5–9). The Icm/Dot translocated substrates (IDTS) hijack host-membrane trafficking pathways, redirecting components of the host cell secretory system to remodel the pathogen compartment into a replication-permissive LCV (10). Most notable is the ability of the LCV to avoid phagosome maturation as a consequence of association with the endoplasmic reticulum (ER), bypassing fusion with compartments that can lead to either microbial degradation or dissolution of the LCV membrane (10–14). Mutations in the Icm/Dot system prevent LCV formation and block intracellular growth, although deletions of single secreted effectors result in either small or undetectable intracellular replication defects. The inability to uncover intracellular growth defects from loss of single effectors is consistent with genetic redundancy, resulting from multiple substrates targeting a single host membrane trafficking pathway or multiple host pathways working in parallel to support LCV biogenesis (6, 15).

The Sde family (*sdeA*, *sdeB*, *sdeC* and *sidE* in the Philadelphia 1 clinical isolate) is a group of homologous IDTS that contains an N-terminal deubiquitinase (DUB), a nucleotidase/phosphohydrolase (NP) domain, and a central mono-ADP-ribosyltransferase domain (mART). The mART domain ADPribosylates host ubiquitin (Ub) that, in turn, is used as a substrate for the NP domain to promote phosphoribosyl-linked Ub (pR-Ub) modification of target host proteins (16–23). One of the primary targets of the Sde family is host reticulon 4 (Rtn4) (19, 22, 24, 25). Phosphoribosyl-Ub modification of Rtn4 promotes endoplasmic reticulum (ER) rearrangements about the LCV within minutes of bacterial contact with host cells (22). Although the absence of Sde family proteins results in small intracellular growth defects in amoebal hosts (26), these defects are subtle during macrophage challenge, consistent with genetic redundancy. This argues that unidentified bacterial translocated effectors may compensate for loss of the Sde family by targeting parallel host pathways that support LCV biogenesis. Redundancy is likely to be limited to the earliest stages of infection, as other IDTS negatively regulate the function of the Sde family. For instance, the mART activity of Sde proteins is inactivated by SidJ, a meta-effector that glutamylates the E860 active site residue (26–29). Furthermore, pR-Ub modification of target proteins is reversed by a pair of pR-Ub-specific deubiquitinases, DupA/B, arguing for temporally limiting Sde family function (24, 25).

In this work, we identified proteins that may compensate for loss of Sde function. Chief among them is the T4SS substrate SdhA protein which is required for maintaining membrane integrity of the LCV (30, 31). SdhA binds the OCRL phosphatase involved in the regulation of early and recycling endosomes, and likely diverts these disruptive compartments from interacting with the LCV (32–34). In the absence of SdhA, the bacteria are exposed to host cytosol and subjected to bacterial degradation by interferon-regulated proteins, leading to pyroptotic host cell death (35, 36). RNAi depletion of Rab5, Rab11 and Rab8, which are guanosine triphosphatases (GTPase) involved in regulating the endocytic and recycling endosome pathways, partially recovers loss of LCV integrity seen in the absence of SdhA, consistent with these compartments disrupting LCV integrity (37).

A second *L. pneumophila* translocated effector that interfaces with the retromer complex is RidL which binds to host VPS29, blocking the function of the retromer which is critical for recycling cargo from endosomes to the trans-Golgi network and to the plasma membrane (38–40). Retrograde trafficking is thought to be blocked by RidL as a consequence of diverting the retromer to sites on the LCV (40), displacing components known to be required for GTP activation of the complex (41).

Using transposon sequencing (Tn-Seq) to unveil redundant effectors involved in LCV biogenesis, we found mutations in three genes (*sdhA*, *ridL* and *legA3*) that aggravate loss of Sde family function. Given the known functions of these effectors, this work argues that Sde proteins act to catalyze formation of a temporally-regulated physical barrier to protect the LCV from attack by host compartments that disrupt the membrane integrity of the replication niche.

## Materials and Methods

### Bacterial strains, cultures, cells and growth media

*L*. *pneumophila* strains were grown in liquid N-(2-acetamido)-2-aminoethanesulfonic acid (ACES) buffered yeast extract (AYE) media or on solid charcoal buffered yeast extract (CYE) media containing 0.4g/l iron (III) nitrate, 0.135 g/ml cysteine, and 1% α-ketoglutaric acid. 40 μg/ml kanamycin, 5% (vol/vol) sucrose, 1 mM IPTG or 5 μg/ml chloramphenicol were added when appropriate. *E*. *coli* strains were cultured in liquid LB or on solid LB plates supplemented with 50 μg/ml kanamycin or 12.5 μg/ml chloramphenicol when appropriate. Primary bone marrow-derived macrophages (BMDM) from AJ mice were prepared and cultured as described previously (13).

### Construction of *L*. *pneumophila* Transposon mutant library

Electrocompetent (42) *L*. *pneumophila* Philadelphia-1 strains SK01(*sde^+^*) and SK02 (D*sde*) were transformed with 75 ng of pTO100*MmeI* (43) respectively, plated on CYE supplemented with kanamycin and sucrose, and incubated at 37°C for 4 days. Multiple pools were made from each strain, each containing 50,000-80,000 colony forming units (CFUs), which were subsequently harvested and pooled into AYE containing 20% (vol/vol) glycerol. Bacterial suspensions were aliquoted at a concentration ~5 × 10^9^ cfu/ml and stored in −80°C. Each of the pools were subjected to deep sequencing of the insertion sites (Illumina HiSeq 2500) to determine pool complexity, and the resulting information was submitted to NCBI Sequence Read Archive under accession No. PRJNA544499.

### Tn-seq screen: Growth of *L*. *pneumophila* Transposon mutant library in BMDM

Three of the transposon mutant library pools in *L. pneumophila* SK01 (*sde^+^*) and SK02 (Δ*sde*) strains, encompassing 186,340 and 170,822 mutants based on deep sequencing analysis, were independently diluted to A600 = 0.25–0.3, cultured to A600 = 0.3–0.4, diluted back to A600 = 0.2 and cultured in AYE to A600 = 3.8-4.0. Aliquots of each library pool grown in AYE were saved as input samples (T1) for growth in AYE broth, and then used to challenge BMDM at a multiplicity of infection (MOI) = 1 for 24 hrs. Cells were washed 3X at 2 hr post-infection (hpi) with PBS, then replenished with fresh medium and further incubated for 24 hrs. BMDMs were then lysed in H_2_O containing 0.05% saponin, and the diluted lysates were incubated on CYE plates with further incubation at 37°C for 3 days. 1-7 × 10^6^ colonies were harvested in AYE, mixed thoroughly and used as the output sample (T2) for sequencing analysis. Genomic DNA from input and output samples was extracted using a QIAGEN DNeasy Blood and Tissue kit, including proteinase K prior to insertion-specific amplification and sequencing.

### Transposon sequencing and Fitness calculation

Illumina^TM^ sequencing libraries were prepared as described previously (44). Genomic DNA (40 ng) was tagmented in a 10 μl reaction mixtures at 55°C for 5 min, followed by inactivation at 95°C for 30s. 40 μl of PCR mixture (First PCR), including primers 1st_TnR, Nextera2A-R and NEB Q5 high-fidelity polymerase (sequences of primers listed in Supplementary Table 1) was added to the tagmented samples to amplify transposon-adjacent DNA. The PCR amplification was performed by incubating at 98°C for 10 s, 65°C for 20 s and 72°C for 1 min (30 cycles), followed by 72°C for 2 min. After amplification, 0.5 μl of the PCR mixture was used in a second PCR reaction containing nested index primers (LEFT indexing primer specific for Mariner and RIGHT indexing primer) and Q5 polymerase in 50 μl total volume. The PCR conditions were 98°C for 10s, 65°C for 20s and 72°C for 1 min, followed by 72°C for 2 min. 9 μl of the PCR reaction was loaded and separated on a 1% agarose-Tris-acetate-EDTA (TAE) gel containing SYBR safe dye, and image intensity in the 250-600 bp region was quantified and pooled from each PCR product in equimolar amounts. The multiplexed libraries were purified on Qiagen QIAquick columns, with 17.5 pmol DNA then used in a 50 μl reconditioning reaction with primers P1 and P2 (Supplemental Table 1) and Q5 polymerase. The reaction was subjected to 95°C for 1 min, 0.1°C/s slow ramp to 64°C for 20 s and 72°C for 10 min. After PCR purification, multiplexed libraries were quantified, size-selected (250-600 bp; Pippin HT) and sequenced (single-end 50 bp) by Tufts University Genomics Core Facility. Sequencing was performed using Illumina HiSeq 2500 with high-output V4 chemistry and custom primer with mar512.

Sequencing reads were processed (FASTX-toolkit), mapped to chromosome (AE017354) (Bowtie) and used to calculate individual transposon mutant fitness using a published pipeline (45). The fitness of an individual mutant (*W*_i_) was calculated based on mutant vs population-wide expansion from T1 to T2 with following equation (46).

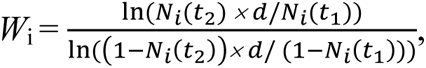

in which *N*_i_(*t*_1_) and *N*_i_(*t*_2_) are the mutant frequency at *t*_1_ and *t*_2_, respectively, and *d* is the population expansion factor. Transposon insertion sites located in the 5’- and 3’-terminal 10% of the open reading frame and genes having less than 5 insertions were excluded for further analysis.

### Construction of *Legionella* deletion mutants

Individual genes were deleted in Lp02 strain by tandem double recombination using the suicide plasmid pSR47s as previously described (47). Primers used to construct all deletion plasmids are listed in Supplementary *SI Appendix* 1, Table S1. Plasmids were propagated in *E. coli* DH5α λpir.

### Construction of complementing plasmids

To perform complementation experiments with genes encoded *in trans* on a replicating plasmid, pMMB207Δ267 (15, 48) was digested with *SacI*-*KpnI* and ligated with PCR-amplified DNA fragments encoding a 6xHis epitope tag, kanamycin resistance gene and *ccdB* flanked by *attR* recombination sites (gift of Tamara O’Connor), which was similarly digested, to generate a Gateway^TM^-compatible destination vector (pSK03; *SI Appendix* 1, Table S1). The individual genes were then cloned into the pSK03 plasmid by integrase cloning from a pDONR221-based IDTS plasmid library (49).

### Data availability

All sequence data are deposited in the NCBI Sequence Read Archive under accession numbers: PRJNA544499, PRJNA847256 and PRJNA864753 (WGS data).

### Code availability

Scripts used for sequencing read analysis can be found at http://github.com/vanOpijnenLab/MaGenTA.

### Intracellular growth assays

The intracellular replication of *L. pneumophila* was measured by luciferase activity using *ahpC::lux* derivatives of the WT and Δ*sde* strains(*SI Appendix* 1, Table S1). Bone marrow-derived macrophages (BMDM) were seeded in 96-well tissue culture plates at a density of 1 x 10^5^ cells per well in RPMI medium without phenol read, containing 10% FBS (vol/vol) and 2 mM of glutamine. BMDMs were incubated at 37°C containing 5% CO_2_ and challenged with *L. pneumophila* Lux^+^ strains at a MOI = 0.05, and luminescence was monitored every 30 min for 3 days during continuous incubation in an environmentally-controlled luminometer (Tecan).

### Quantitative RT-PCR

PMA-differentiated U937 cells were seeded at a density of 4 x 10^6^ cells per well in 6 well tissue culture plates. Cells were infected with post-exponentially grown *L. pneumophila* Lp02 at a MOI = 20 and washed at 1 hpi. RNA was isolated using Trizol in accordance with manufacturer’s instructions. Contaminating genomic DNA was removed using the TURBO DNA-free kit, and cDNA was synthesized with 2.5 μg of RNA using SuperScript VILO cDNA Synthesis Kit. PowerUp SYBR Green Master Mix was then used for qRT-PCR reactions using Second Step instrument (ABI). Transcriptional levels of genes were normalized to 16S rRNA. Oligonucleotides are listed in *SI Appendix* 1, Table S1.

### Cytotoxicity assays

10^5^ BMDMs were seeded per well in 96 well tissue culture plates and incubated overnight at 37°C, 5% CO_2_ prior to replenishing with 100 μl of medium containing propidium iodide (PI) at a final concentration = 20 μg/ ml. Cells were challenged with 100 μl of postexponentially grown *L. pneumophila* Lp02 in the same medium at a MOI = 1 or 5 (36). Following infections, cells were centrifuged at 400 x g for 5 mins, incubated at 37°C, 5% CO_2_, and PI uptake was monitored every 10 min using the bottom reading setting in an environmentally controlled fluorometer (Tecan). To determine 100% cytotoxicity as the normalization control, cells were treated with 0.1% Triton X-100, and PI uptake was determined.

### Assay for vacuole integrity

To measure the fraction of infected cells having intact *Legionella-*containing vacuoles (LCV), bacteria were centrifuged onto BMDMs for 5 min and incubated at 37°C, 5% CO_2_ for noted periods of time. The infection mixtures were then fixed in PBS containing 4% paraformaldehyde, then probed with mouse anti-*L. pneumophila* (Bio-Rad, Cat# 5625-0066, 1:10,000) followed by secondary probing with goat anti-mouse Alexa Fluor 594 (Invitrogen, Cat# A11005, 1:500), to identify permeable vacuoles as described (31). After washing 3X in PBS, LCVs were permeabilized by 5 min incubation with −20°C methanol, prior to a second probing with mouse anti-*L. pneumophila*. All bacteria (both from intact and disrupted vacuoles) were identified by goat anti-mouse IgG Alexa Fluor 488 (Invitrogen, Cat# A11001, 1:500). The amount of vacuole disruption was quantified in two fashions. First, individual BMDMs were imaged using Zeiss observer Z1at 63X and scored for permeabilization based on staining with goat anti-mouse IgG-AlexaFluor 594 (antibody accessible in absence of methanol permeabilization) as described previously (32). To allow larger numbers of infected cells to be imaged, automated microscopy was performed using the Lionheart FX scanning microscope and Gen5 image prime 3.10 software. For detection of disrupted vacuoles (permeable in absence of methanol treatment), all images were analyzed by image preprocessing (10X magnification). To determine colocalization and quantification of vacuole integrity at 2 hpi, a primary mask was set for goat anti-mouse IgG-AlexaFluor 488 (detected after methanol treatment) and a secondary mask was set using a region that was expanded approximately 0.001 μm from the primary mask for goat anti-mouse IgG Alexa Fluor 594 (detected before methanol treatment). To identify intracellular bacteria, DAPI staining of nuclei was used to threshold a secondary mask 4 μm apart from the primary mask.

### RTN4 colocalization with LCV

RTN4 colocalization with the LCV was assayed by immunofluorescence microscopy. BMDMs were infected with *Legionella* strains for 4 hrs, fixed in PBS containing 4% paraformaldehyde, then extracted in 5% SDS to remove most of the cell-associated RTN4, then probed with mouse anti-*L. pneumophila* (Bio-Rad, Cat# 5625-0066, 1:10,000) and rabbit anti-RTN4 (Lifespan Biosciences, Cat# LS-B6516, 1:500) to detect detergent-resistant structures about the LCV. Bacteria were detected with anti-mouse Alexa fluor-594 and RTN4 structures with anti-rabbit Alexa Fluor-488 (Jackson ImmunoResearch, Cat# 711-545-152, 1:250).

### Nucleofection

Differentiated BMDMs were seeded at a density of 5 x 10^6^ cells in 10 cm dishes filled with 10 mls RPMI medium containing 10% FBS and 10% supernatant produced by 3T3-macrophage colony stimulating factor (mCSF) cells (50) and incubated overnight. Cells were lifted in cold PBS and resuspended in RPMI medium containing 10% FBS. Resuspended cells were aliquoted into 1.5 ml microfuge tubes containing 1 x 10^6^ cells and pelleted at 200 x g for 10 min. The pellets were resuspended in nucleofector buffer (Amaxa Mouse Macrophage Nucleofector Kit, Cat# VPA-1009) and 2 μg of siRNA was added (siGENOME smart pool, Dharmacon). Cells were transferred to a cuvette and nucleofected in the Nucleofector 2b Device using Y-001 program settings according to manufacturer’s instructions. Nucleofected macrophages were immediately recovered in the medium and plated in 8-well chamber slides at 5 x 10^4^ /well for microscopy assays or in 12-well plates at 1.5 x 10^5^/well to prepare cell extracts for immunoblotting.

### Immunoblotting

The efficiency of siRNA silencing in nucleofected cells was determined by immunoblot probing of SDS-PAGE fractionated proteins. Necleofected macrophages plated in 12-well plates were lysed by incubating in RIPA buffer (Thermo Fisher Scientific, Cat#89900) for 20 min on ice and protein concentration was measured by BCA assay. 5-10 μg of protein in SDS-PAGE sample buffer was boiled for 10 min, fractionated by SDS-PAGE and transferred to nitrocellulose membranes. The membrane was blocked in 50 mM Tris-buffered saline/0.05% Tween 20 (TBST, pH 8.0) containing 4% nonfat milk (blocking buffer) for 1 hr at room temperature and probed with primary antibodies against Rab5B (Proteintech, Cat# 27403-1-AP, 1:1,000), SNX1 (Proteintech, Cat# 10304-1-AP, 1:1000), RTN4 (Lifespan Biosciences, Cat# LS-B6516, 1:2,000), polyHistidine (Sigma-Aldrich, Cat# H1029, 1:2,000) and β-actin (Invitrogen, Cat# PA1-183, 1:1,000) in blocking buffer at 4°C overnight. After washing 3X with TBST, the membranes were incubated with secondary antibody (Li-Cor Biosciences, Cat#926-32211, 1: 20,000) in blocking buffer for 45 min at room temperature. Capture and analysis were performed using Odyssey Scanner and the image Studio software (LI-COR Biosciences).

## Results

### Identification of genes involved in LCV biogenesis that can compensate for the loss of Sde

We previously demonstrated that the *L. pneumophila* Δ*sde* strain (Δ*sidE* Δ*sdeABC*) is partially defective for growth within protozoan hosts, but grows in murine macrophages at levels close to that of the *L. pneumophila* WT strain (22, 51). This phenomenon is consistent with the existence of redundant Icm/Dot translocated substrates (IDTS) that can compensate for the lack of Sde proteins in mammalian hosts (15). To identify redundant pathways involved in intracellular growth, we performed transposon sequencing (Tn-seq) mutagenesis to uncover mutations that aggravate the D*sde* intracellular growth defect within bone marrow-derived macrophages (BMDMs). Transposon library pools were generated in both the WT and the Δ*sde* strain using the *Himar1* transposon which specifically inserts at TA dinucleotides (52).

Three independently collected pools of *Himar-1* insertions were constructed in the *L. pneumophila* WT and D*sde* strains, encompassing 117,419 (47.33% of total TA sites) and 108,934 (43.91% of total TA sites) total unique insertions in the two genomes, respectively. This represented approximately 34 and 31 insertions/gene in WT and D*sde*, respectively (Dataset S1). After growth in broth to post-exponential phase (T1) (53), BMDMs were challenged with both pools for 24 hr (equivalent to a single round of infection; T2) and the fitness contribution of each mutation was determined during growth in broth and in BMDMs (Fig. 1A; Materials and Methods) (46). To identify genes that were required for intracellular replication in macrophages, the fitness difference between BMDM growth versus nutrient-rich medium growth was calculated in the WT and D*sde* strains, respectively (Figs. 1B, C). Insertions in the majority of genes that were nonessential for growth in broth exhibited a fitness of ~1 during 24 hr incubation in BMDMs, indicating that most genes are not required for intracellular growth in macrophages (*SI Appendix* 1, Fig. S1). It has been established that individual loss of only 6 IDTS (*mavN*, *sdhA*, *ravY*, Lpg2505, *legA3* and *lidA*) impair growth in macrophages, while individual loss of most of the other 300+ effectors show little defect in intracellular growth. This has been attributed to functional redundancy in *Legionella* secreted effectors (10, 30, 54–57). In our datasets, we confirmed those genes were required for replication in macrophages in WT (Fig. 1B, indicated by blue lettering). Furthermore, mutations in the preponderance of genes encoding translocator effectors generated no statistically significant defects in intracellular growth in either of the two backgrounds (Figs. 1B, C), consistent with previous studies.

**Fig. 1.**
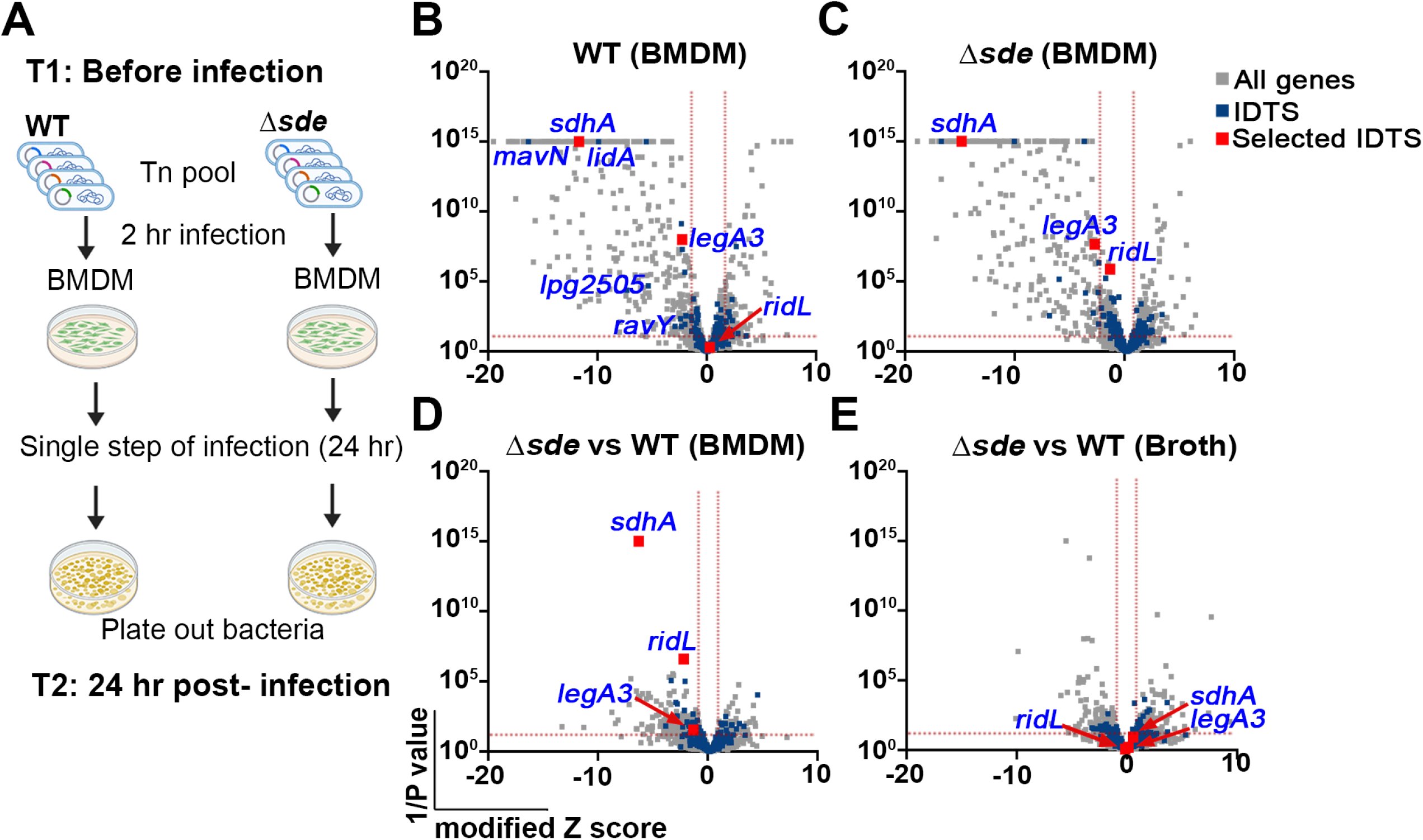
Tn-seq identifies mutations that aggravate loss of Sde function. (**A**) Schematic view of Tn-seq analysis to identify aggravating mutations. *Himar-1* pools were constructed in parallel in SK01 (WT) and SK02 (Δ*sde*) strains and insertion site abundance was determined after growth in broth (Materials and Methods). Three of the sequenced pools were incubated with BMDMs for 24 hrs in parallel, plated on bacteriological medium, and relative abundance of insertions was determined by HTS to determine fitness of individual mutations in the two different strain backgrounds (Materials and Methods). (**B, C**) Volcano plots of the relative fitness, represented as modified Z scores (Materials and Methods) comparing replication in BMDM versus AYE for either WT or Δ*sde* strains. Candidates were identified based on criteria of Z_MOD_ > 2 from the population median and were statistically significant (p < 0.05) based on unpaired t-test after Two-stage step-up correction (indicated by dotted red line) (73, 74). Blue font indicates IDTS that are the focus of study or which were previously shown to have an intracellular growth defect. (D) Volcano plots displaying relative fitness of insertion mutations in a Δ*sde* background compared to the WT background for intracellular growth in BMDM. Genes were identified as candidates based > 1 MAD from the population mean and statistical significance (p < 0.05) based on unpaired t-test after Two-stage step-up correction method (74) (indicated by dotted line). Data are based on n=3 biological replicates of pools made in each strain. (E) Volcano plots of relative fitness (modified Z scores) of mutations in Δ*sde* background versus WT background for growth in AYE broth culture. Grey, blue and red squares represent whole genes, *icm/dot* or Icm/Dot translocated substrate (IDTS) genes and genes selected based on the following criteria, respectively. Criteria: 1) mutations who showed fitness differences (Δ*sde* - WT) > 1 median absolute deviation (MAD) from the population median fitness and were statistically significant based on unpaired t tests (p < 0.05; Dataset S1), 2) genes that were *Icm/Dot* translocated substrates, and 3) genes possibly involved in LCV biogenesis.

We then filtered mutations based on the following criteria: 1) causing lowered fitness relative to the population median, as defined by modified Z score > 1 (median absolute deviation (MAD) >1) and p< 0.05 based on unpaired t tests comparing mutations in the WT vs. Δ*sde* background (Dataset S1); 2) genes encoding *Icm/Dot* translocated substrates; and 3) genes encoding proteins thought to be involved in LCV biogenesis, based on published data. In addition, we identified mutations that showed no statistical defect in the WT, but whose fitness difference (Δ*sde* - WT) was > 1 MAD from the population median without any other consideration. Based on these criteria, 3 genes (*sdhA*, *ridL* and *legA3*) were prioritized for further analysis (Fig. 1D). SdhA protein is a T4SS substrate required for maintaining LCV integrity (30, 31). The absence of *sdhA* caused a growth defect in BMDMs infected with either WT or the Δ*sde* strains (Fig. 1B, C). Even so, the fitness defect was significantly aggravated when *sdhA* was disrupted in the Δ*sde* strains relative to its loss in a WT strain background (Fig. 1D).

Mutations in *ridL* or *legA3* also showed aggravating growth defects in the D*sde* strains (Fig. 1D). RidL is a T4SS substrate that can inhibit function of the retromer complex that modulates retrograde traffic from early endosomes (40, 58). LegA3 is an ankyrin-repeat effector protein that is required for optimal replication in several hosts that has not been clearly tied to replication vacuole formation previously (15, 38, 43). Defective growth was specific to BMDMs as the double mutant strains grew as well as the WT strain in bacteriological medium (Fig. 1E and Dataset S1). Most notably, based on their efficient growth in a WT background, was the behavior of insertions in *ridL*, which showed clear defects in a D*sde* background (Figs. 1C, D)

### The absence of Sde proteins exacerbates intracellular growth defects of *sdhA*, *ridL* or *legA3* deletion mutants

To verify that the phenotypes predicted by the parallel Tn-seq pools can be reproduced at the single strain level, in-frame deletion mutations in *sdhA, ridL and legA3* were generated in both the WT and D*sde* strain backgrounds harboring the luciferase (*ahpC*::*lux^+^*) reporter. The respective mutations were confirmed by whole genome sequencing (PRJNA864753), and intracellular growth of *L. pneumophila* strains was then monitored by luminescence accumulation after incubation with BMDMs. As predicted by the Tn-seq analysis, combining the loss of Sde proteins with the D*ridL* mutation revealed a growth defect that did not exist in the absence of the combination. In the presence of the Sde proteins, the D*ridL* strain showed no growth defect after challenge of BMDMs. In contrast, introduction of D*ridL* into the D*sde* strain resulted in yields that were 100X lower relative to the WT and approximately 10X lower relative to the parental D*sde* strain after 72 hr incubation (Fig. 2A). Additionally, catastrophic synergistic defects were observed with the D*legA3* mutation after introduction into the D*sde* background. In an otherwise WT background, loss of either Sde proteins or RidL resulted in mild growth defects after 72 hours incubation. The D*sde*D*legA3* strain, however, showed little or no evidence of growth during this time period (Fig. 2B). Finally, although the D*sdhA* strain reproduced the previously documented growth defect in BMDMs (30, 31), combination with D*sde* resulted in a strain that showed yields similar to the type IV secretion system defective *dotA^−^* strain (Fig. 2C)

**Fig. 2.**
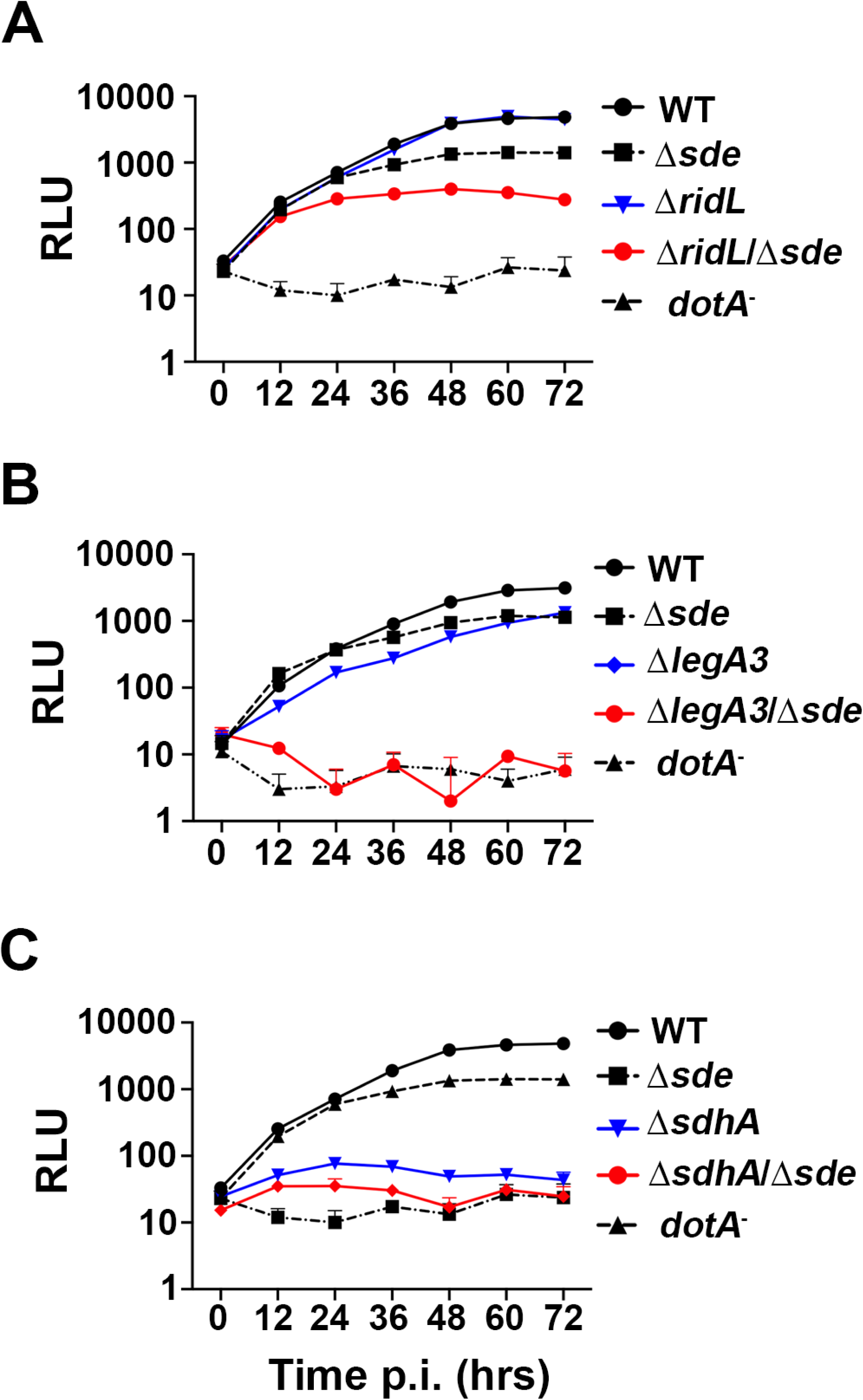
Identification of mutations that aggravate the intracellular growth defect of Δ*sde*. (A-C) BMDMs were challenged with WT (Lp02) or noted *L. pneumophila* mutants expressing luciferase (P*ahpC*::lux). Intracellular growth was determined by measuring luminescence hourly. Data shown and error bars are mean ± SEM at 12 hr increments (mean of 3 technical replicates and a representative of 3 biological replicates).

### Aggravation of the D*sde* lesion causes premature host cell death due to destabilization of the LCV

To determine if there were a clear defect in replication vacuole biogenesis associated with aggravating the loss of Sde function, we took advantage of the fact that SdhA is required to maintain integrity of the LCV, reasoning that the absence of Sde could exacerbate this defect (31, 32). As had been noted previously, challenge with a D*sdhA* strain resulted in a significant fraction of the LCVs becoming permeable and accessible to antibody penetration at 6 hours post-infection (hpi) (*SI Appendix* 1, Fig. S2) (31). At 2 hpi, however, there is no evidence that the absence of SdhA interferes with LCV integrity, with both the WT and D*sdhA* strains showing indistinguishable levels of permeability to antibody staining (*SI Appendix* 1, Fig. S2). As Sde proteins act to remodel Rtn4 about the replication vacuole within 10 min of bacterial challenge (31, 51), we reasoned that any compensation for loss of SdhA should occur at early timepoints. Therefore, we sought to examine the integrity of the LCV in BMDMs at 2 hpi.

Vacuole integrity of the mutants was evaluated by probing fixed BMDM with anti-*L. pneumophila* in the presence or absence of chemical permeabilization, using our previously established immunofluorescence staining method (Fig. 3A) (31). Surprisingly, even the Δ*sde* single mutant strain generated a higher frequency of permeable vacuoles than WT at the 2 hr timepoint (Fig. 3B). In contrast, *ridL*, *legA3* and even *sdhA* single deletions showed vacuole permeability frequencies that were comparable to WT at this timepoint (Fig. 3B). The most dramatic effects were observed when deletions of *sdhA*, *ridL*, and *legA3* were introduced into the Δ*sde* strain. Addition of each individual deletion to the Δ*sde* mutant severely aggravated the vacuole integrity defect, resulting in up to 4-fold more permeable vacuoles, indicating that these mutation combinations drastically destabilized the LCV (Fig. 3B). To allow larger numbers of LCVs to be analyzed (1000 to 3000 per biological replicate), albeit at lower resolution, we used a lower power objective to repeat the analysis. Although low power analysis made it more difficult to identify permeable vacuoles, the results were in concordance with those displayed in Fig. 3B (*SI Appendix* 1, Fig. S3). These results indicate that at early timepoints after infection, the *L. pneumophila* has redundant T4SS substrates that can compensate for the loss of Sde, and that vacuole disruption resulting from absence of *sde* is potentiated by the addition of secondary mutations in *sdhA*, *ridL* and *legA3*.

**Fig. 3.**
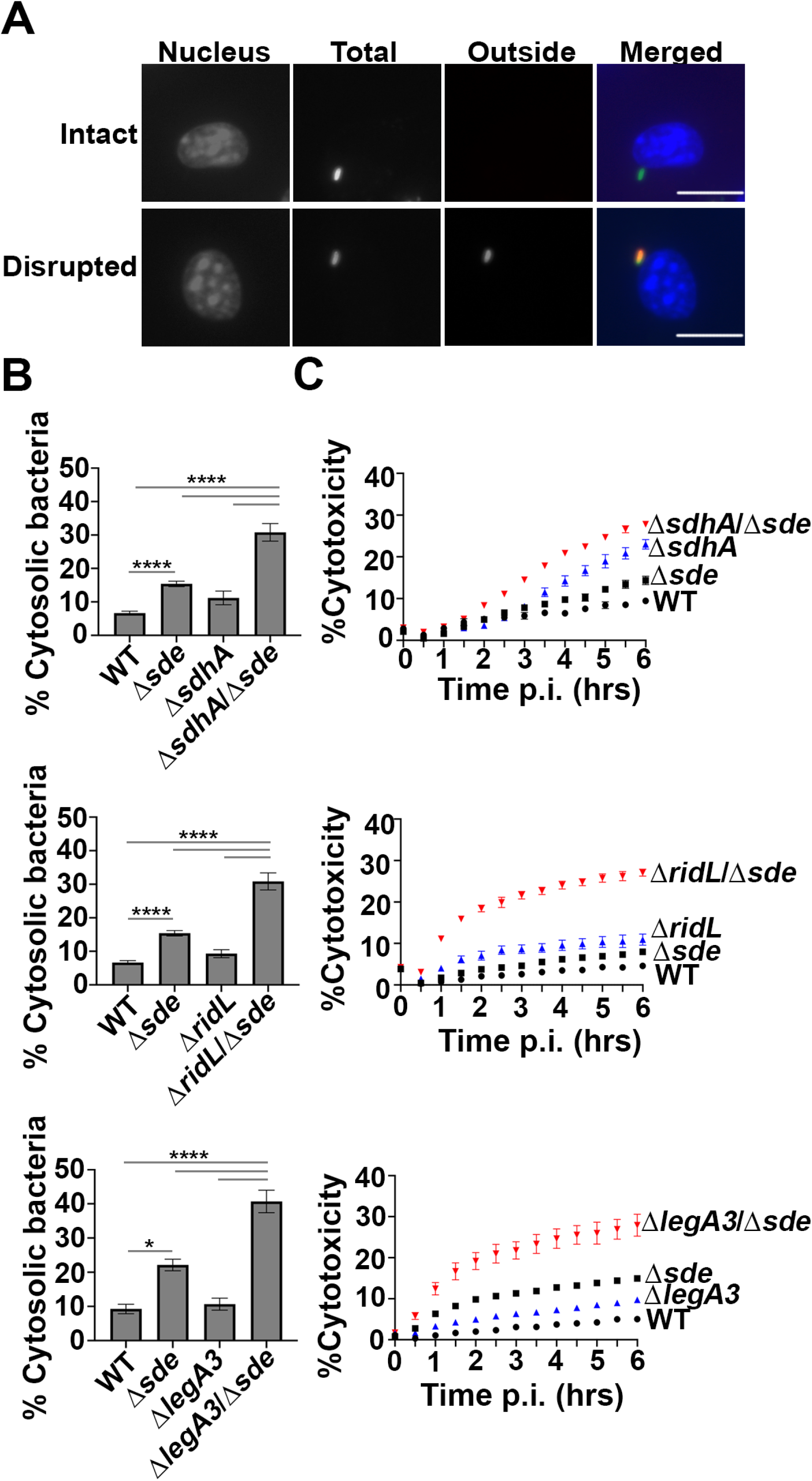
Aggravating mutations result in loss of LCV integrity and accelerated pyroptotic cell death at early infection times. (A) Examples of cytosolic and vacuolar bacteria. Macrophages were challenged with either WT or noted mutant strains for 2 hr, fixed, probed with anti-*L. pneumophila* (Alexa Fluor 594 secondary, red), permeabilized, and reprobed with anti-*L. pneumophila* (Alexa Fluor 488 secondary, green). Cytosolic bacteria are accessible to both antibodies, shown in yellow in the merged image, whereas vacuolar bacteria are shown in green. The scale bar represents 10 µm. (B) Disrupted vacuole integrity of *L. pneumophila* strains at 2 hr post-infection. BMDMs were challenged with indicated strains, fixed and stained for bacteria before and after permeabilization. For quantification, bacteria that stained positively in the absence of permeabilization were divided by the total infected population (mean ± SEM; three biological replicates with 300 LCVs were counted per replicate) (C) Kinetics of macrophage cell death, with infection of indicated strains. BMDMs were infected with *L. pneumophila* WT or mutant strains and propidium iodide (PI) incorporation was used to monitor cell death. Data shown and error bars are mean ± SEM for 30 min increments and a representative of 3 biological replicates. Statistical analysis was performed using one-way ANOVA with Tukey’s multiple comparisons, with significance represented as: *p<0.05; ****p < 0.0001.

Pyroptotic cell death occurs as a consequence of a comprised LCV membrane followed by bacterial exposure to the macrophage cytosol (31). Based on the loss of LCV barrier function shortly after initiation of infection, we hypothesized that the respective Δ*sdhA*Δ*sde*, Δ*legA3*Δ*sde* and Δ*ridL*Δ*sde* double mutant strains should accelerate BMDM cell death in comparison to either the WT or single mutants. To this end, cytotoxicity assays were performed, assaying for propidium iodide (PI) access to the macrophage nucleus (Materials and Methods; (36)). In perfect concordance with the loss of LCV integrity, pyroptotic cell death was exacerbated when mutations in either *sdhA*, *ridL* and *legA3* were combined with D*sde* (Fig. 3C). Accessibility to PI was more rapid than that observed with the WT and the total cytotoxicity plateaued at levels that far exceeded the WT during the course of the experiment, consistent with the early defect of these double mutants resulting in increased overall damage to the LCV relative to the WT.

### Downmodulation of Sde activity by SidJ interferes with vacuole protection

To demonstrate that expression of single Sde effectors was sufficient to restore LCV integrity in the absence of SdhA function, plasmids encoding individual effectors were introduced into the D*sde*D*sdhA* strain. As expected, the introduction of plasmid-encoded *sdhA* resulted in increased LCV stability, as measured by the antibody accessibility assay (Fig. 4A). Plasmids encoding single Sde effectors (*sdeA*, *sdeB*, *sdeC*: Fig. 4A) allowed similar levels of LCV protection to that observed for the *sdhA*-harboring plasmid. We conclude that each of the Sde proteins demonstrated previously to catalyze phosphoribosyl-linked ubiquitination of Rtn4 and promote associated Rtn4 rearrangements (Fig. 4B; (22)) is sufficient to allow protection of the LCV from degradation.

**Fig. 4.**
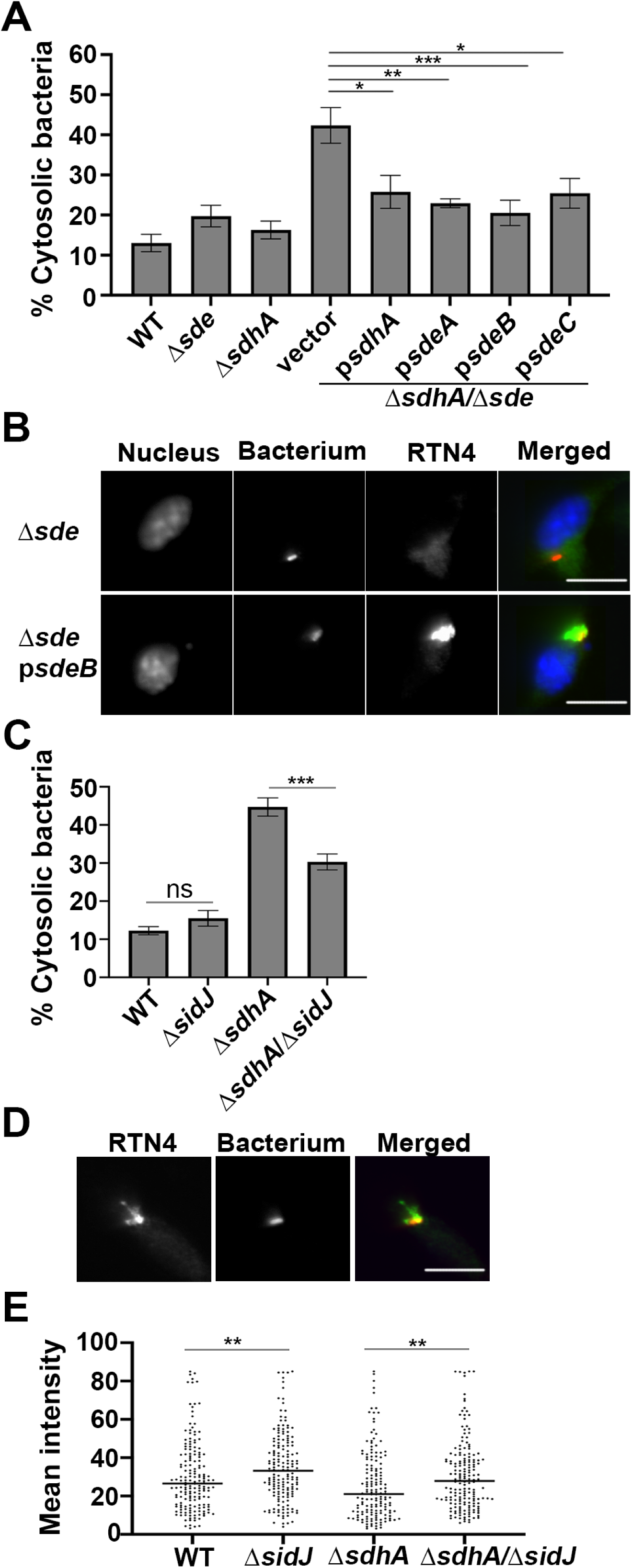
Increased aggregation of RTN4 is associated with maintenance of LCV integrity. (A) Vacuole integrity of *L. pneumophila* strains at 2 hr post-infection. BMDMs were challenged with indicated strains, fixed and stained for bacteria before and after permeabilization. The percentages of cytosolic bacteria were calculated. Data displayed as mean ± SEM for three biological replicates. (B) Complementation of Δ*sdhA*Δ*sde* strain *in trans*. Shown is aggregation of RTN4 in fixed samples of BMDM challenged with *L. pneumophila* Δ*sdhA*Δ*sde* harboring p*sdeA*. Bacteria and RTN4 were probed as described (Materials and Methods), scale bar 10 µm. (C) Vacuole integrity of *L. pneumophila* strains at 4 hr post-infection. Data shown as mean ± SEM for four biological replicates. At least 70 LCVs were counted per replicate (A, C). Statistical significance was tested using one-way ANOVA with Tukey’s multiple comparisons; *p<0.05,**p<0.01, ***p<0.001 (A, C). (D) A Representative micrograph of RTN4 associated with individual LCVs, scale bar 10 µm. (E) Quantification of RTN4 intensity associated with individual LCVs. Loss of SidJ results in increased association of RTN4 about LCV. Images of individual LCVs were captured and pixel intensities of RTN4 staining about regions of interest were determined. More than 70 LCVs were quantified per experiment and data were pooled from 3 biological replicates (**p<0.01; Mann-Whitney test)

Sde proteins are only able to compensate for loss of SdhA at early times after infection of BMDMs as there was an accumulation of degraded LCVs in the D*sdhA* strain even in presence of Sde at later time points (*SI Appendix* 1, Fig. S2; (31)). An explanation for this phenomenon is that protection of the LCV by Sde proteins is negatively regulated in a temporal fashion by the SidJ meta-effector which shuts down Sde activity by glutamylation of the mART domain (E860 residue on SdeA; (28)). The primary consequence of this post-translational modification is that Sde proteins are released from the LCV (26), but active SidJ shutdown of phosphoribosyl-linked ubiquitination is also predicted to prevent accumulation of proteins such as Rtn4 about the LCVs. Therefore, the absence of SidJ should both prolong Sde activity and act to compensate for loss of SdhA as the infection proceeds beyond the 4 hr timepoint. To test this hypothesis, a deletion of *sidJ* was introduced into the Δ*sdhA* strain and vacuole integrity was examined at 4 hpi (Fig. 4C). Based on the antibody protection assay, at 4 hpi there was ~33% reduction in degraded vacuoles as a consequence of removing SidJ from the Δ*sdhA* strain (Fig. 4C; compare Δ*sidJ*Δ*sdhA* to Δ*sdhA*).

Deletion of SidJ did not completely protect from vacuole disruption, as there were clearly more intact vacuoles after infection with SdhA^+^ strains, indicating that pR-Ub-linked targets likely persist for extended periods of time in the presence of intact SidJ function (WT compared to Δ*sidJ*Δ*sdhA*; Fig. 4C). One likely candidate for persistent modification is RTN4 (Fig. 4D), which forms detergent-resistant aggregates that may slowly dissipate in the absence of continued modification by Sde proteins. To determine if restoration of vacuole disruption is connected to partial loss of RTN4 aggregates, the intensity of Rtn4 aggregates around the LCV was quantified microscopically after detergent extraction (Materials and Methods). Consistent with expectations, the total RTN4 intensity (area plus unit intensity) was increased by removing SidJ (Fig. 4E). The effect was independent of the presence of the *sdhA*, as in both WT (Fig. 4E; WT vs Δ*sidJ*) and Δ*sdhA* strains (Fig. 4E; Δ*sdhA* vs. Δ*sidJ*Δ*sdhA*) the absence of SidJ results in higher levels of Rtn4 accumulation at 4 hpi. Therefore, loss of SidJ lengthens the time that Sde function can compensate for loss of SdhA and is associated with increased accumulation of Rtn4 at this timepoint.

### LCVs harboring strains lacking *sde* are stabilized by depletion of proteins involved in early endocytic and retrograde trafficking

Based on work from both *Salmonella* and *Legionella*, the most common explanation for bacterial mutants that result in destabilization of the pathogen replication vacuole is that there is a failure to protect from association with host membrane compartments that result in vacuole degradation (37, 59). For example, SdhA antagonizes function of the early endocytic compartment, in part by diverting the OCRL phosphatase (32, 37). As a consequence, depletion of components that regulate early endocytic dynamics, such as Rab5bB, prevents LCV degradation, stabilizing the replication niche in the absence of SdhA (32, 37, 60, 61). To determine if Sde and SdhA proteins protect the LCV from attack by the same host membrane trafficking pathways, depletion with small interfering RNA (siRNA) against Rab5B in BMDMs was performed and vacuole integrity was measured at 2 hpi. At this timepoint, when Sde proteins appear to be the primary stabilizers of the LCV, knockdown of Rab5B partially reversed vacuole disruption after challenge with the Δ*sdhA*Δ*sde* strain (Fig. 5A). This indicates that Sde and SdhA are likely able to interfere with the same early endocytic pathway. As expected, no restoration was observed in cells infected with WT, although the corresponding single Δ*sde* strain, which shows a small defect in maintaining vacuole integrity, was unaffected by the loss of Rab5B (Fig. 5A). Therefore, in the absence of Rab5B, there may be another pathway that Sde proteins directly antagonize to promote vacuole integrity.

**Fig. 5.**
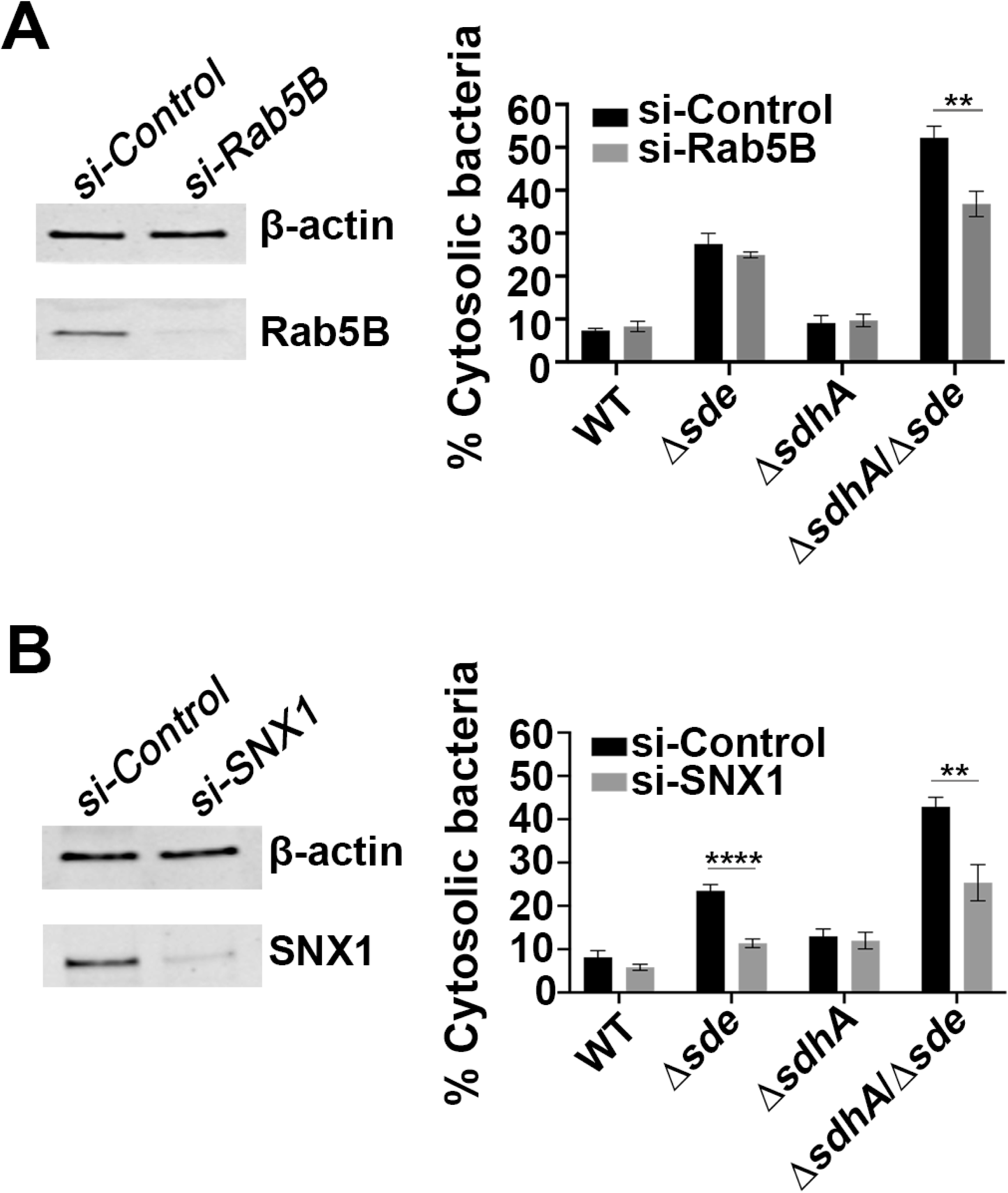
Interference with host endocytic and retromer-mediated trafficking pathways allows partial rescue of vacuole integrity defect. (A) Effect of si-Rab5B on vacuole integrity. Left: immunoblot of SDS-polyacrylamide gel probed with anti-Rab5B after depletion with si-Rab5B or control scrambled si-RNA. Right: the percentage of cytosolic bacteria from BMDMs treated with si-Control or si-Rab5B, and challenged with noted *L. pneumophila* strains for 2 hrs, followed by probing as described (Materials and Methods). (B) Effect of si-SNX1 on vacuole integrity. Left: immunoblot of SDS-polyacrylamide gel probed with anti-SNX1 after depletion with si-SNX1 or control scrambled si-RNA. Right: the percentage of cytosolic bacteria from BMDMs treated with si-Control or si-SNX1 and challenged with noted *L. pneumophila* strains for 2 hrs, followed by probing as described (Materials and Methods). For immunoblotting, β-actin was used as loading control. The data shown are mean ± SEM, using three biological replicates, with at least 100 LCVs counted per biological replicate. Statistical analysis was conducted using unpaired two-tailed Student’s t test and significance displayed as **p < 0.01 or ****p < 0.0001.

We previously demonstrated that knockdown of Rab11, a GTPase that regulates the recycling endosome, partially rescues the vacuole integrity defect of a Δ*sdhA* strain. This indicates that host recycling compartments interfere with vacuole integrity (37). Furthermore, Sorting Nexin 1 (SNX1) participates in these events, and functions in tandem with the Retromer, which is a target of RidL (39, 40, 62). We thus predicted that the depletion of SNX1 should also reverse vacuole disruption. To this end, SNX1 was depleted by siRNA prior to bacterial infection and vacuole integrity was measured at 2 hpi. In this case, the ability to partially rescue the vacuole integrity defect at an early timepoint was independent of *sdhA* function. For both the Δ*sde* and Δ*sde*Δ*sdhA* strain backgrounds, depletion of SNX1 resulted in a significant reduction of disrupted vacuoles, with 50% and 38% fewer antibody-permeable vacuoles observed in the presence of siRNA directed toward SNX, respectively, when compared to treatment with the scrambled control (Fig. 5B). These results support previous work that early endosome dynamics are involved in disrupting the LCV and are consistent with Sde proteins playing a special role in blocking SNX1/retromer-mediated membrane traffic at early timepoints after bacterial challenge of primary macrophages.

### Temporally regulated IDTS maintain LCV integrity during infection

Sde proteins act immediately after contact of bacteria with mammalian cells to promote Rtn4 rearrangements and maintain LCV integrity (22). In contrast, SdhA acts when Sde activity appears to dissipate due to continued action of SidJ (Fig. 4; (26–29), as a Δ*sdhA* strain requires four hours to exhibit a clear defect in LCV integrity. Therefore, these proteins likely execute their roles sequentially, with Sde proteins functioning as vacuole guards at the earliest stages of replication compartment formation, while SdhA works downstream at later time points. To support this sequential function, we hypothesized that there should be temporal regulation of the vacuole guards identified here. To this end, we measured the relative transcription of the vacuole guards after bacterial contact of a WT strain with cultured cells. The expression of *sdeA* and *sdeC* was high at 1 hpi and then dramatically decreased by about 10-fold at 6 hpi compared to that at 1 hpi (Fig. 6). Expression of SidJ was maintained, or raised gradually throughout the infection, consistent with its role in downmodulating Sde function. As found previously, the level of *ridL* expression gradually receded from 1 to 6 hpi (Fig. 6; (40)). In contrast, the expression of *sdhA* and *legA3* increased during infection, by 4- or 10-fold respectively, at 6 hpi (Fig. 6A). These results are consistent with SdhA and LegA3 playing critical roles in preserving LCV integrity at infection times that extend beyond the initial establishment of the LCV (~4 hr) (31).

**Fig. 6.**
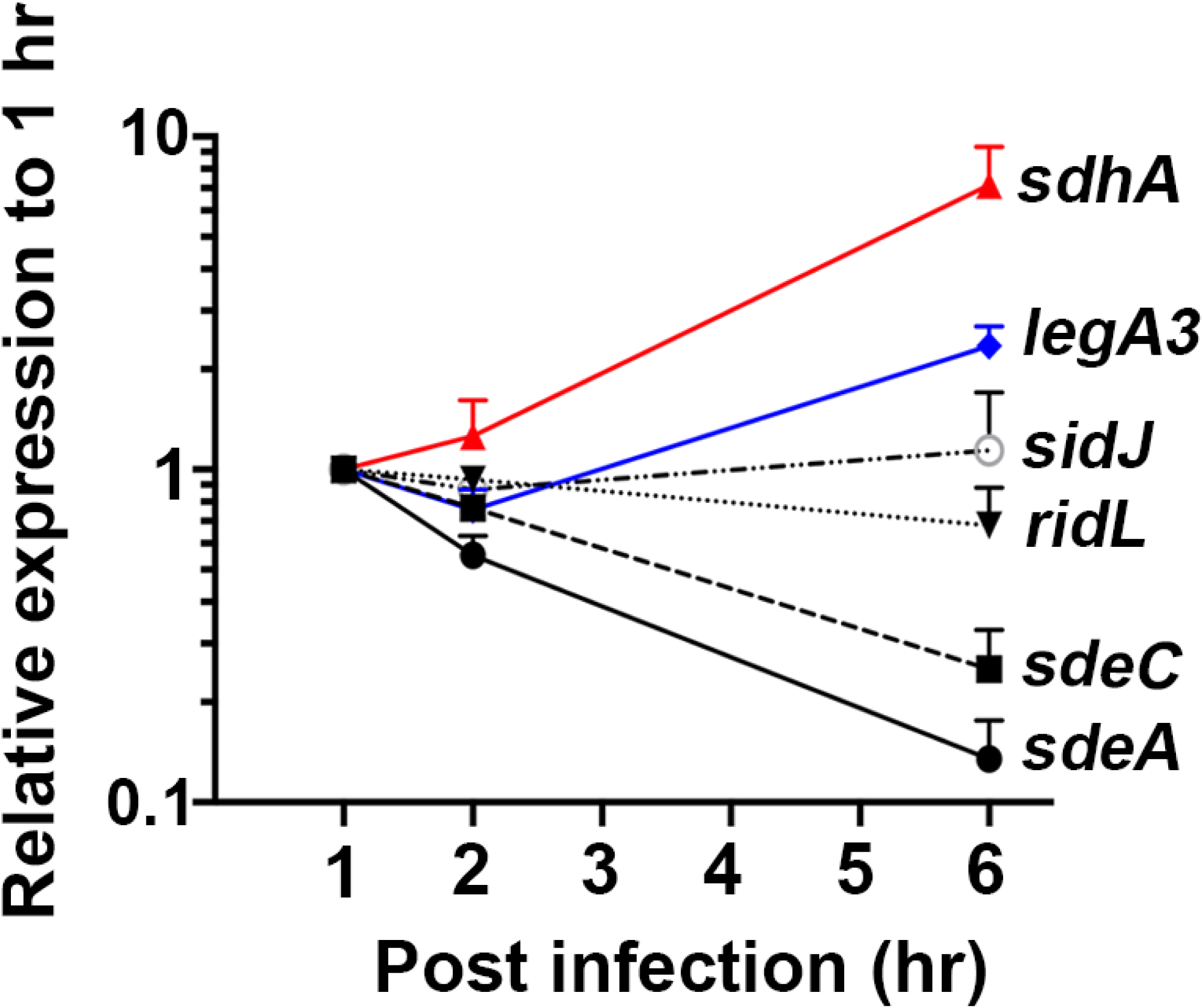
Transcription of *sde* genes is downregulated during *L. pneumophila* infection of BMDMs. Transcript abundance of indicated genes was determined during infection. PMA-differentiated U937 cells were challenged with *L. pneumophila* WT and RNA was extracted at the noted time points. Transcripts were normalized to 16s rRNA, and then displayed relative to transcription level measured at 1hr post-infection and represented as fold change.

## Discussion

Replication vacuole integrity is a critical determinant of successful pathogen growth within membrane-bound compartments (63). The importance of this process has been demonstrated for several intracellular pathogens, all of which encode proteins critical for maintaining an intact vacuolar barrier (31, 64–66). In each case, bacterial proteins appear to interfere with the function of membranes exiting from early endosomal/recycling compartments (32, 37, 40). There is no clear explanation for how endosomal membranes disrupt replication compartments, but bacterial mutant studies indicate that the replication vacuole has a unique membrane composition that is destabilized by endosomal membranes (67). In particular, *S. typhimurium* and *L. pneumophila* mutants lacking specific phospholipases stabilize these compartments (31, 68). In the case of *L. pneumophila plaA* mutations, loss of a lysophospholipase reduces the fraction of permeable LCVs observed in *sdhA* mutants, indicating that modulation of lysophospholipid content may maintain replication vacuole integrity. In fact, analysis of the *L. pneumophila* translocated effector VpdC argues that lysophospholipid content regulates LCV expansion, consistent with vacuole integrity being dependent on homeostatic control of lysophospholipids (69).

In this report, we obtained the surprising result that the *L. pneumophila* Sde proteins contribute to maintaining LCV integrity (Fig. 2), consistent with their blocking endosomal membrane traffic (Fig. 5). Loss of SdhA or RidL aggravated the minor growth defect of a Δ*sde* strain, thereby eliminating proteins that interface directly with endosomal factors (Fig. 1). In the case of SdhA, the protein engages and diverts the OCRL phosphatase endosomal traffic regulator, while RidL prevents activation of retromer components that modulate retrograde traffic from endosomes (32, 40, 41, 58). Sde proteins may work in a very different fashion than these two proteins. Sde proteins catalyze phosphoribosyl-linked modification of a large number of host proteins, modifying Ser and perhaps Tyr residues on protein targets (19, 70). An early target is the endoplasmic reticulum protein Rtn4, which is not known to directly interface with the endosomal system (22). Largescale identifications of proteins modified by SdeA confirm that Rtn4 is among the most abundant pR-UB-linked proteins (24, 25).

The connection between ER rearrangements and support of membrane integrity raises the possibility that Sde action protects the LCV from host attack by forming a shield about the LCV. Immediately after *L. pneumophila* contact with mammalian cells, Sde proteins drive abundant accumulation of Rtn4-rich tubular ER aggregates (Fig. 7). These structures result in complex pseudovesicular structures, observed over 20 years ago in both mammalian cells and amoebae within 10 minutes post-infection (14, 22, 71). Over time, these structures dissipate and are replaced by rough ER (14, 71). Therefore, regulation of Sde function ensures that it operates primarily during the initial stages of replication. This explains why loss of SdhA has little effect on LCV integrity during the first 2 hpi (Fig. 3), and only shows a defect when combined with loss of Sde function. Therefore, SdhA primarily plays a backup role in the first 2 hpi (Fig. 7), but when Rtn4-rich pseudovesicular structures dissipate at later time points (22, 51), SdhA assumes its role as the primary essential guard against LCV disruption.

**Fig. 7.**
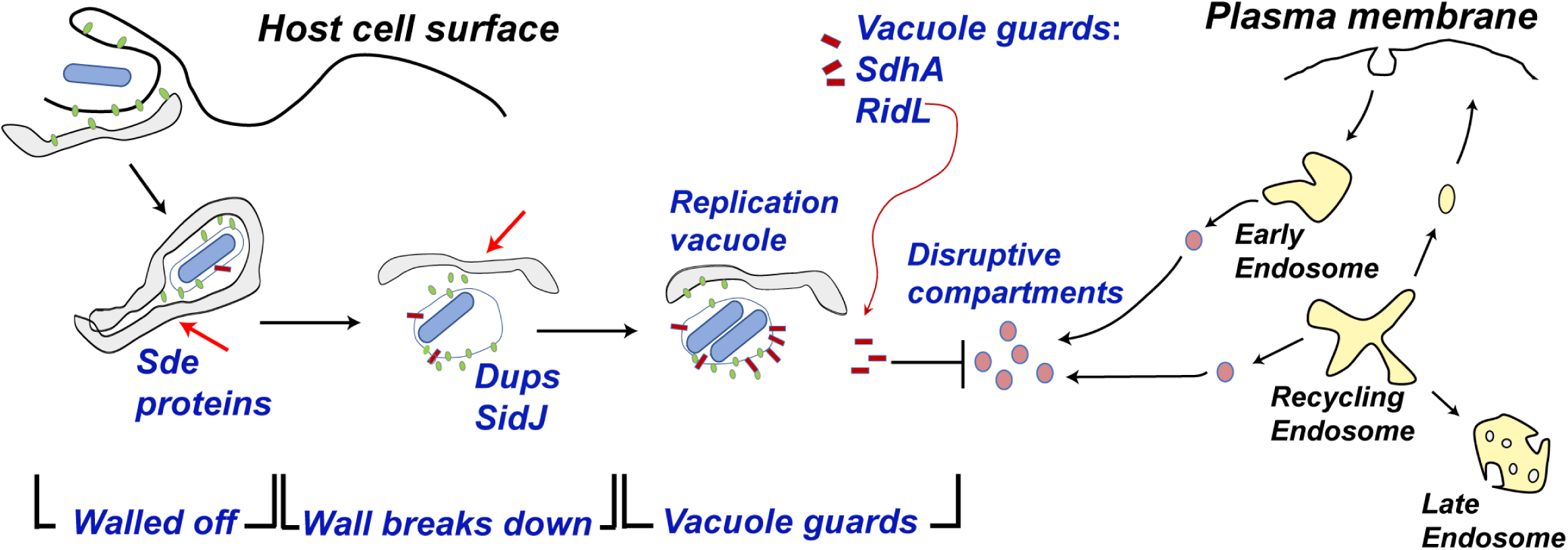
Schematic model of how *Legionella* combats vacuole disruption in a temporally-regulated process. Soon after internalization, the *Legionella*-containing vacuole is walled off with RTN4-rich tubular ER aggregates as a consequence of PR ubiquitination on RTN4 by Sde proteins (22) **[Step 1. Walled off]**. At early time of infection, the barrier as well as SdhA and RidL (backup vacuole guards in case the barrier is breached) protects the LCV from host membrane traffic derived from the early endosome. Over time, the wall is breached as the aggregates are dissipated by SidJ, a metaeffector that inactivates Sde proteins, (26–29) and Dups (DupA and DupB), enzymes that deubiquitinate PR-Ub-linked substrates (24, 25) **[Step 2. Wall Breaks Down]**. When the barrier around the LCV is dismantled, SdhA and RidL act to divert and inactivate disruptive compartments derived from the early/recycling endosomes to allow the LCV to be replication-permissive (32, 37) **[Step 3. Vacuole guards]**.

Based on these results, we propose a model in which Sde, RidL and SdhA promote LCV integrity in a temporally controlled fashion (Fig. 7). Shortly after infection, we hypothesize that the Sde proteins act to wall off the LCV from endosomal attack by rearranging tubular ER into dense structures as a consequence of pR-UB-modification of Rtn4 (22). RidL and SdhA are present on the LCV to protect against occasional breaches of this barrier at early time points (32, 40). The continued presence of the physical barrier, however, is likely to pose problems for supporting bacterial growth because it prevents access of the LCV to either metabolites or lipid biosynthesis components. This predicts that optimal growth of *L. pneumophila* must involve the breakdown of the pR-Ub-modified physical barrier. Therefore, metaeffectors that reverse Sde family function, such as the SidJ protein (27–29), are necessary for optimal intracellular growth because they facilitate barrier breakdown (26, 72). Consistent with this model, there is suboptimal growth in the absence of SidJ (26, 72), with the negative consequence that *L. pneumophila sidJ^−^* mutants accumulate Rtn4 at the 6 hr timepoint (Fig. 5). To avoid a tradeoff between supporting vacuole integrity and interfering with intracellular growth, *L. pneumophila* has acquired the ability to disrupt the Sde-promoted barrier, necessitating the localization of the SdhA vacuole guard on the LCV to protect a newly established point of vulnerability for the replication niche (Fig. 7).

This work provides a fresh view of the role of redundancy in an intracellular pathogen. In the case of protecting LCV integrity, our work argues that multiple proteins do not work in parallel pathways toward the same end. Instead, each pathway is temporally controlled, playing an important role at different times in the replication process. This then explains the profound replication defect of a *sdhA^−^*strain, which only has some low-level support from RidL and residual remnants of the Sde-targeted protein blockade. The lack of effective backup pathways as the replication cycle proceeds necessitates the up-regulation of SdhA, resulting in a largely nonredundant role for this protein (Fig. 6). That RidL and LegA3 are not particularly effective backups for SdhA as the infection cycle proceeds, indicates that these proteins may be unable to block critical membrane-disruptive pathways that are inactivated by SdhA.

An important caveat to this model is that Sde protein can target a number of proteins other than Rtn4 (16, 24, 25). Furthermore, Sde localization is not restricted to the LCV, but family members can be found on a number of organelles, including endosomes/lysosomes and mitochondria at early stages of infection (~ 1 hr post-infection) (26). In this regard, we think it likely that there are two modes of action that can promote vacuole integrity. One mode is to establish a physical barrier around the LCV by eliciting Rtn4-ER rearrangements. The other is to modify host proteins associated with endocytic trafficking, Golgi biogenesis or autophagy (25). In this regard, partial rescue of the LCV integrity defect by depletion of SNX1 in BMDMs infected with Δ*sde* mutants is particularly noteworthy (Fig. 5C). Previous studies have shown that SNX1 is localized on LCVs, raising the possibility that proteins controlling the movement, docking and fusion of disruptive compartments could come in contact with Sde proteins and allow inactivation of these compartments (25, 40). Therefore, Sde proteins may act as their own backup factors, inactivating disruptive compartments that sneak through the Rtn4-aggregated barrier.

In summary, by performing parallel dense transposon mutagenesis in matched strains, we have obtained evidence that the Sde family acts to protect the LCV from disruption by the host. Furthermore, our study provides a new framework for vacuole guard function, as the described guards are temporally regulated to maximize the replication potential of *L. pneumophila*. Future work will focus on how manipulation of host membrane compartments leads to maintaining LCV integrity, and determining the molecular details for how LCV disruption occurs in the absence of vacuole guards.

## Acknowledgements

We thank Dr. Joseph Vogel for the kind gift of the *Legionella* strain (Δ*sidJ*), Drs. Ila Anand and Mengyun Zhang for preliminary work indicating that SNX1 contributes to driving LCV disruption, and Dr. Philipp Aurass for discussions regarding analysis of Tn-seq results. We thank Juan Hernandez-Bird, Mitchell Berg, and Drs. Wenwen Huo, Philipp Aurass and Kevin Manera for review of the text. This work was supported by NIAID Awards 5R01-AI46245 and 5R01-AI113211 to RRI.

## Supporting Information

**Fig. S1.**
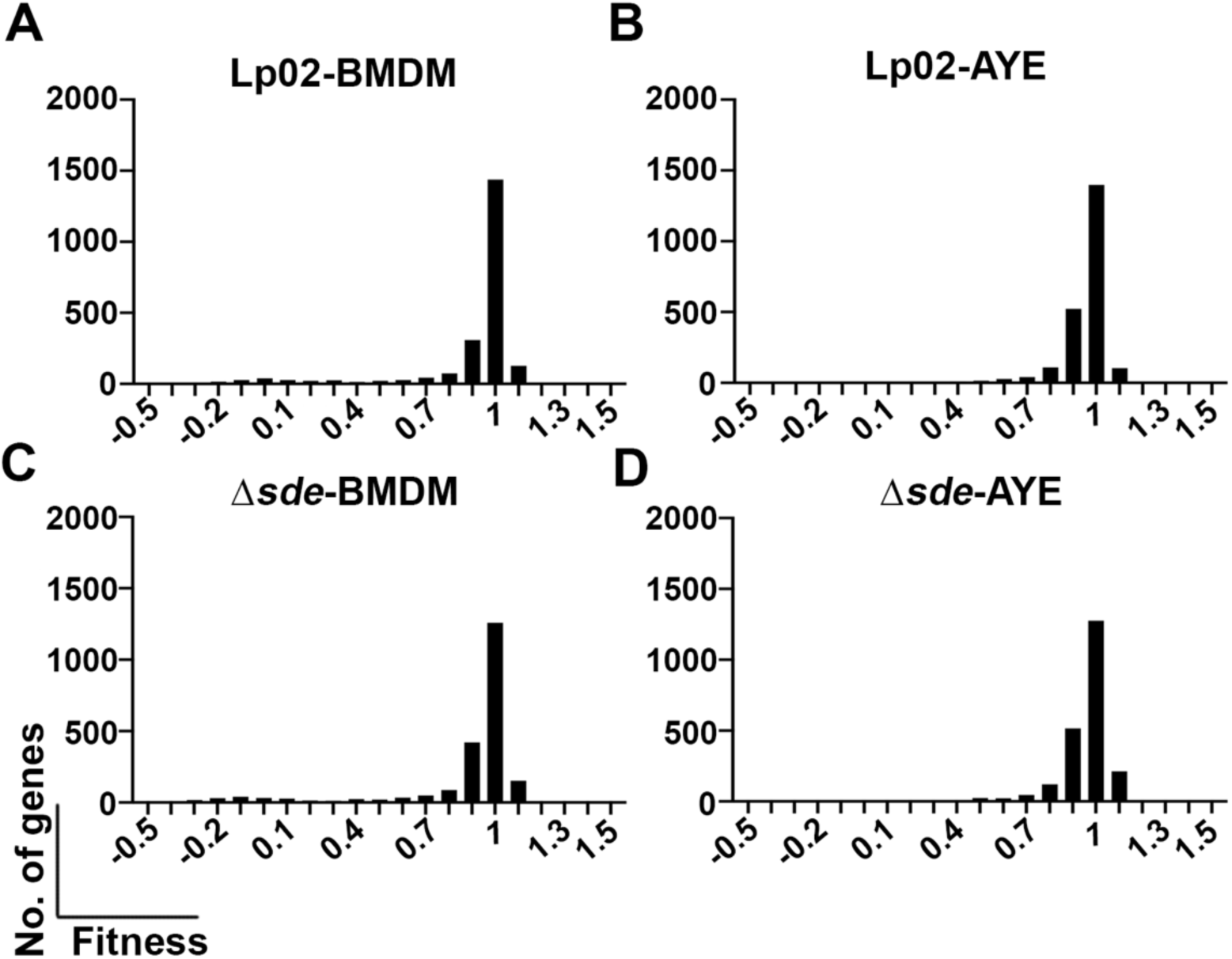
(Linked to Fig.1). Histogram plots of fitness for all *L. pneumophila* genes represented on Tn-seq. Histogram of WT (SK01) Tn-seq pool following either infection in BMDM (A) or growth in nutrient-rich AYE medium (B). Histogram of Δ*sde* (SK02) Tn-seq pool following infection in BMDM (C) or growth in nutrient-rich AYE medium (D).

**Fig. S2.**
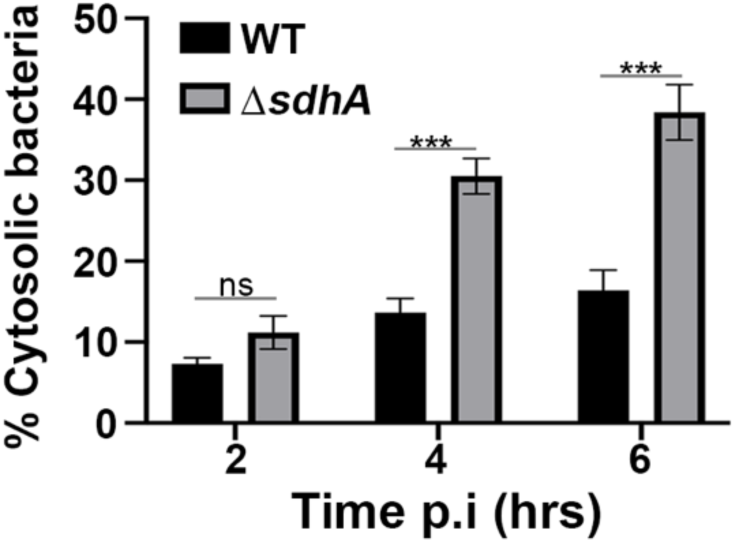
(Linked to Fig.3). The integrity of LCVs harboring Δ*sdhA* strains after challenge with *L. pneumophila*. Percent cytosolic bacteria was quantified based on antibody accessibility. BMDMs were infected with either WT or Δ*sdhA* strains for 2, 4, and 6 hr, fixed, and stained with antibodies. The internalized bacteria in the absence of permeabilization were calculated relative to the total infected population (mean ± SEM; three biological replicates were performed and 100 LCVs were counted per biological replicate). Statistical analysis was conducted using unpaired two-tailed Student’s t test (ns, not significant; ***p < 0.001).

**Fig. S3.**
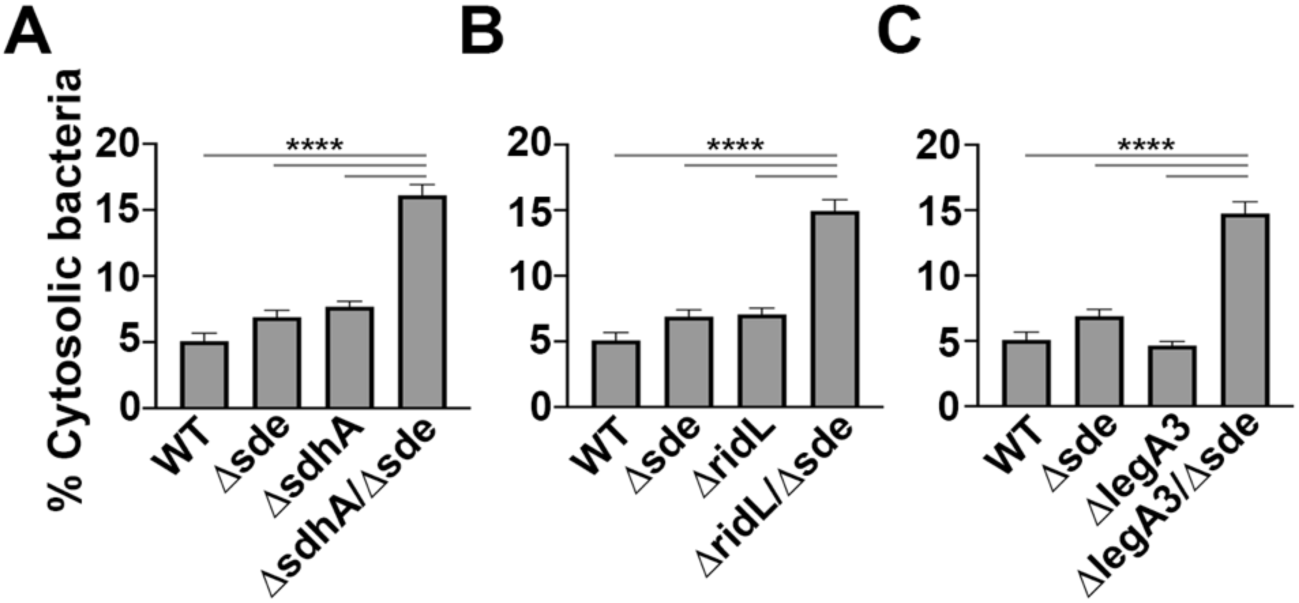
(Linked to Fig. 3). The loss of *sdhA*, *ridL* and *legA3* aggravated vacuole disruption in 11*sde* strain. Vacuole integrity was measured based on antibody accessibility. BMDMs in a 96 well plate were infected with the indicated strains for 2 hr, fixed and stained with antibodies. The images were taken by Lionheart automatic microscope using 10X magnification objective. The internalized bacteria in the absence of permeabilization were calculated relative to total infected population to determine fraction of disrupted vacuoles (mean ± SEM; three biological replicates were performed and 1000-3000 LCVs were counted per biological replicate). Statistical significance was tested using one-way ANOVA with Tukey’s multiple comparisons; ***p <0.001.

**Table S1.**
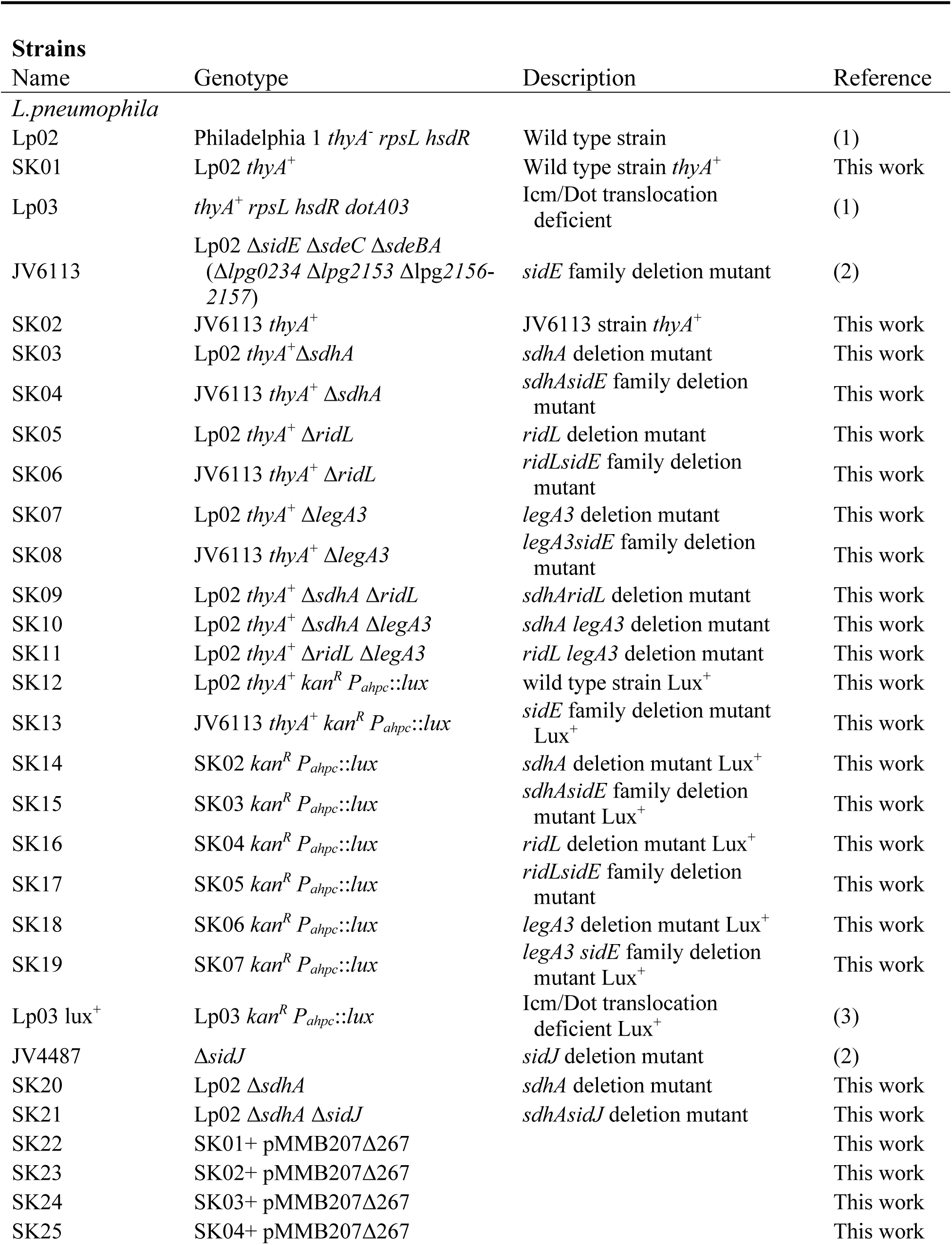
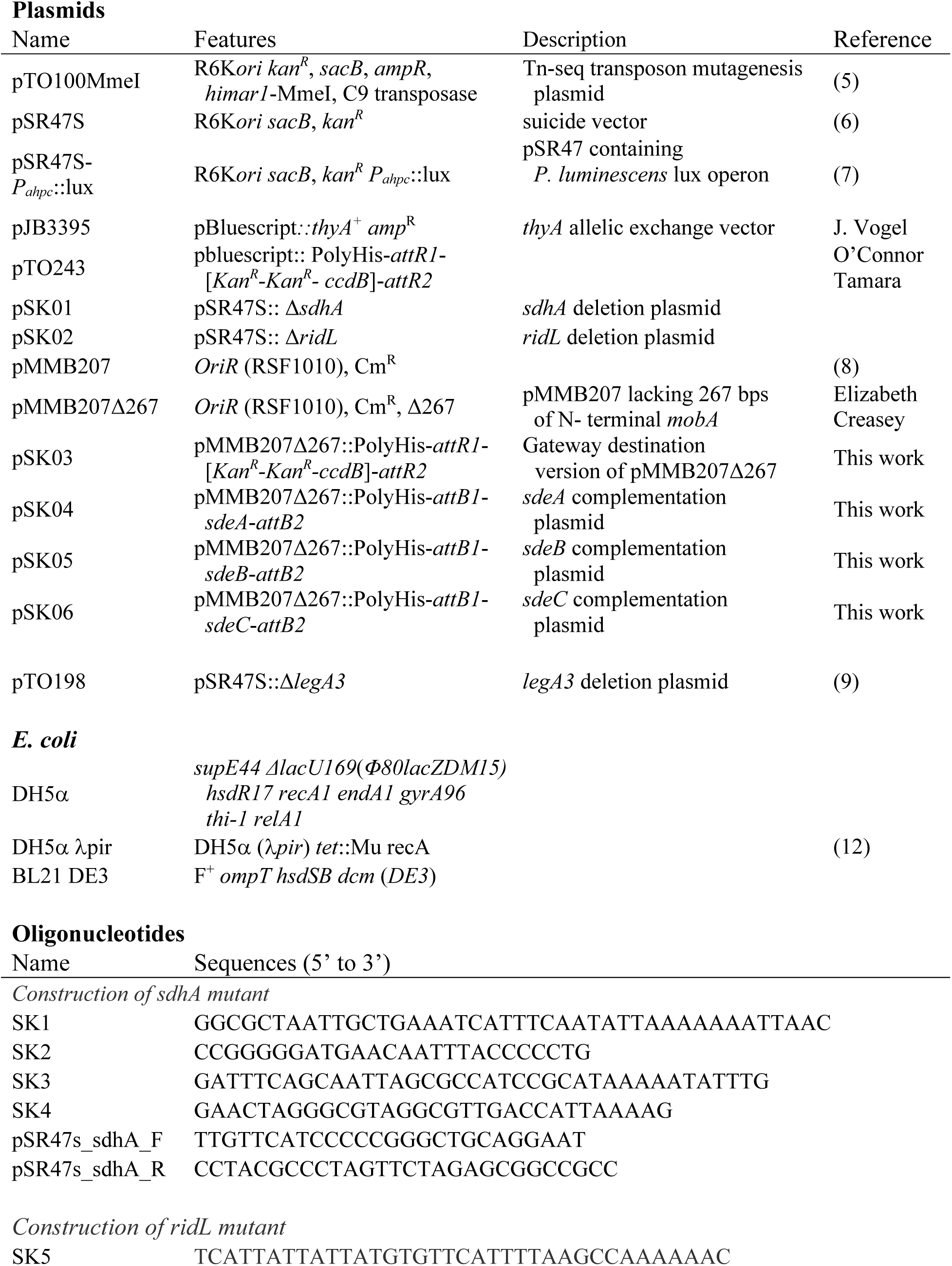
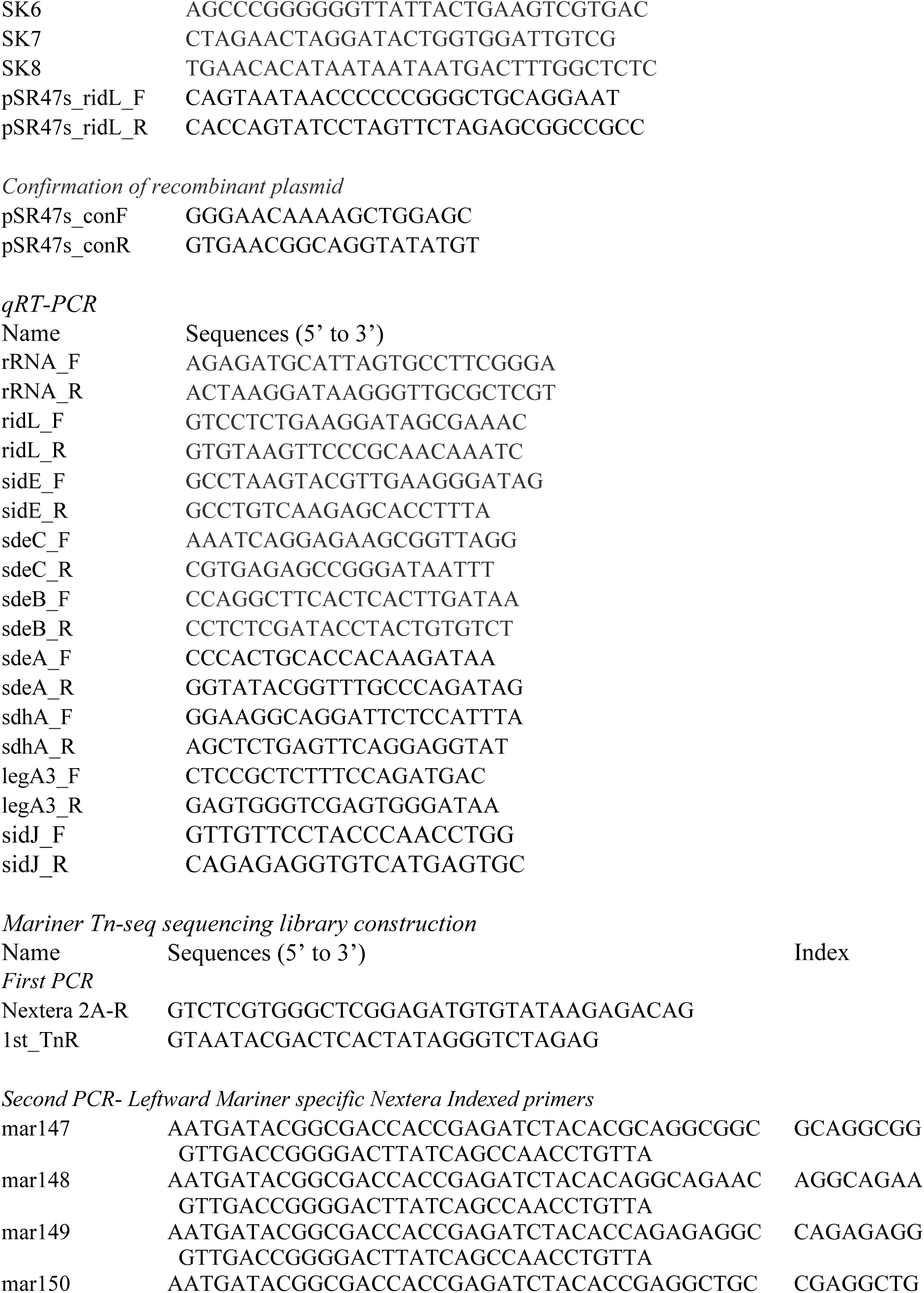
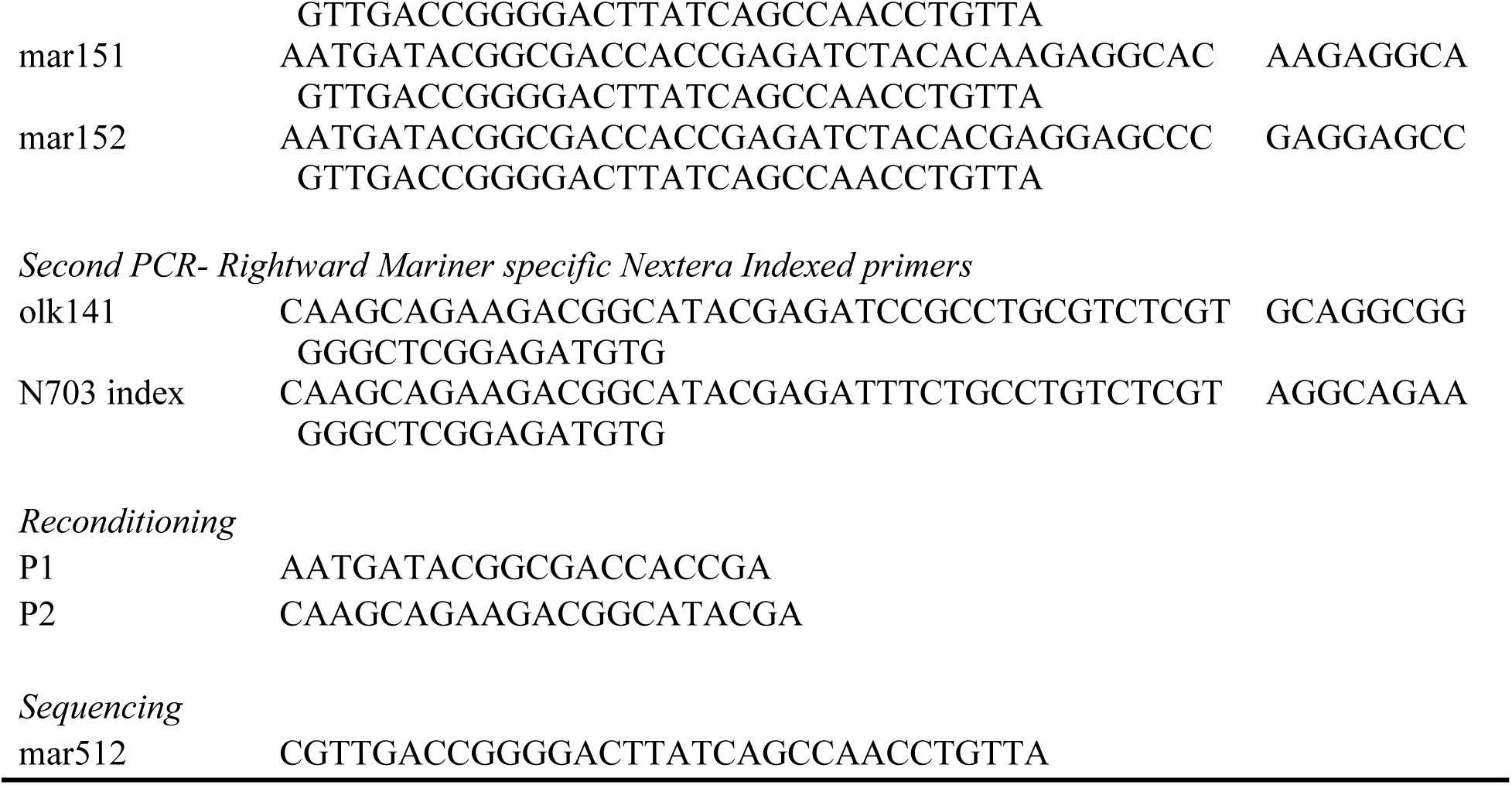
Strains, Plasmids and Oligonucleotides used in this study.

